# Somatic mutation but not aneuploidy differentiates lung cancer in never-smokers and smokers

**DOI:** 10.1101/2023.01.05.522947

**Authors:** Sitapriya Moorthi, Amy Paguirigan, Minjeong Ko, Mary Pettinger, Anna C. H. Hoge, Anwesha Nag, Neil A. Patel, Feinan Wu, Cassie Sather, Matthew P. Fitzgibbon, Aaron R. Thorner, Garnet L. Anderson, Gavin Ha, Alice H. Berger

## Abstract

Lung cancer in never-smokers disproportionately affects older women. To understand the mutational landscape of this cohort, we performed detailed genome characterization of 73 lung adenocarcinomas from participants of the Women’s Health Initiative (WHI). We find enrichment of *EGFR* mutations in never-/light-smokers and *KRAS* mutations in heavy smokers as expected, but we also show that the specific variants of these genes differ by smoking status, with important therapeutic implications. Mutational signature analysis revealed signatures of clock, APOBEC, and DNA repair deficiency in never-/light-smokers; however, the mutational load of these signatures did not differ significantly from those found in smokers. Last, tumors from both smokers and never-/light-smokers shared copy number subtypes, with no significant differences in aneuploidy. Thus, the genomic landscape of lung cancer in never-/light-smokers and smokers is predominantly differentiated by somatic mutations and not copy number alterations.

## Introduction

Lung cancer is the deadliest cancer in both men and women worldwide, with lung adenocarcinoma being the most prevalent subtype^1^. The clinical management of lung adenocarcinoma has transformed dramatically over the last decade owing to the discovery of driver oncogenes such as *EGFR* and the development of corresponding targeted therapies^2^. These discoveries have been pivotal in clinically changing the course of lung cancer. However, much of the genetic characterization of lung cancer has been performed on tumors from patients with a history of smoking^3,4^. Cigarette smoking is the primary risk factor for developing lung cancer. Yet 10-15% of lung cancer cases in the U.S., and up to 20% of cases worldwide, occur in patients who have never smoked, defined as individuals who have smoked <100 cigarettes^5–7^. In recent years, the percentage of lung cancer cases in ‘never-smokers’ has increased to 17% in men and 24% in women, which may reflect both a decrease in global smoking behavior and an increase in the incidence of lung cancer in never-smokers ^8^. A particular concern is that the incidence of lung cancer in young women appears higher than in young men, even after controlling for differences in smoking behavior^9^. If considered as a separate disease, lung cancer in ‘never-smokers’ would be the 7th largest cause of death due to cancer^5,10^. Thus, even as smoking rates decline, lung cancer in never-smokers is expected to contribute to a significant cancer burden in the U.S. and worldwide.

Lung cancer in never-smokers is distinct from lung cancer in smokers due to many unique genetic and clinical characteristics^11,12^. The most frequently diagnosed histological subtype of lung cancer in never-smokers is adenocarcinoma^5^, women are diagnosed more often than men^13,14^, and the majority of lung cancer cases in south and east Asian women occur in never-smokers^5^. Although lung cancer in never-smokers affects young adults^15^, older individuals are at a higher risk of diagnosis and constitute the majority of never-smokers lung cancer patients in the U.S.^16^.

At the genetic level, lung tumors from smokers have a significantly higher overall somatic mutation rate and different somatic mutation patterns than tumors from never-smokers, suggesting alternative mechanisms of cancer development in never-smokers and smokers^4,17^. Tumors from never-smokers are enriched for *EGFR* mutations and fusions involving *ALK, RET, ROS1*, or *NRG1*^*18– 20*^, and have less frequent *KRAS* mutations than tumors from smokers^21^. Recently, the NCI SHERLOCK study reported whole genome sequencing of 232 lung cancers from never-smokers^11^. The authors described the clustering of tumors from these never-smokers into distinct groups based on arm-level copy number alterations, which correlated with prognosis. However, because these subtypes were defined primarily from tumors from never-smokers, it remains unclear if copy number subtypes are unique to never-smokers or if these subtypes are present in lung cancer in general.

Here, we sought to define the genetic landscape of lung cancer in female never-smokers along with a matched cohort of smokers. We performed genomic analysis of tumor and matched normal DNA from post-menopausal Women’s Health Initiative (WHI) participants who developed lung cancer during the study^22^. We find that never-smokers display a unique mutational spectrum of *EGFR* and *KRAS* variants with implications for both targeted and immunotherapy. Moreover, we surprisingly did not detect chromosomal fusions in *ALK, RET*, and *ROS1* suggesting that lung cancers from older female never-smokers may have lower rates of these fusion oncogenes. Somatic mutation signature analysis found DNA repair defect signatures in 22% of the tumors, although we were unable to attribute this phenotype to germline cancer predisposition variants. Finally, we confirm the recent finding of distinct copy number subtypes of lung adenocarcinoma^11^, but we find that these subtypes are shared across tumors from smokers and never-smokers. Thus, we find that aneuploidy and somatic copy number alteration are general features of lung cancer not related to smoking.

## Results

### Genomic profiling of lung cancer in female never-smokers

Lung cancer in never-smokers occurs in more older women compared to younger women **(Figure S1A)**^16^, and unique clinical and molecular characteristics distinguish younger and older cancer patients^15,23,24^. To understand the genomic landscape of lung cancer in older never-smokers women we identified post-menopausal women from the WHI LILAC study^25,26^, who developed lung adenocarcinoma **(Table S1)**. The WHI was initially conceived as three overlapping clinical trials and an observational study to evaluate risk factors for cancer and cardiovascular disease^22,26^. The study cohort comprised women with either less than 100-lifetime cigarettes (‘never-smokers) or a light-smoking history of fewer than five pack-years. A smaller group of heavy smokers with greater than 20 pack-year smoking history were matched to this cohort on cancer stage, diagnosis year, and tumor purity. To enable the detection of both exonic mutations and known chromosomal fusions, we performed custom whole exome sequencing on matched tumor and normal DNA samples (**Methods**) (tumor median target coverage 93x; tumor mean target coverage range 13.95x-214.58x; normal median target coverage 80x; normal mean target coverage range 45.68x-122.41x). In total, 73 tumor-normal pairs from 56 never-/light-smokers and 17 heavy smokers passed the quality control assessment and were used for downstream analysis (**Table S2 and S3)**.

### Profound differences in somatically mutated genes in tumors from smokers and never-smokers

The single nucleotide variant (SNV) and insertions-deletions (indel) landscapes of never-/light-smokers showed extensive differences (**Figure 1 and Table S4**). Tumors from heavy smokers had a higher non-silent tumor mutational burden (TMB) and a greater percent of C-to-A transversion mutations compared to tumors from never-smokers **(Figure 1A-B)**, consistent with cigarette smoke being a direct mutagen of the genome^3,4,17^. Moreover, smoke exposure, measured by the pack-years of cigarette smoked, significantly correlated with both TMB (r^2^ = 0.3758) and percent of C-to-A transversions (r^2^ = 0.3905) (**Figure 1C-D**). Tumors from never- and light-smokers had indistinguishable TMB, and C-to-A mutation rates (**Figure 1A and C**), and thus were grouped for subsequent analyses.

**Figure 1.**
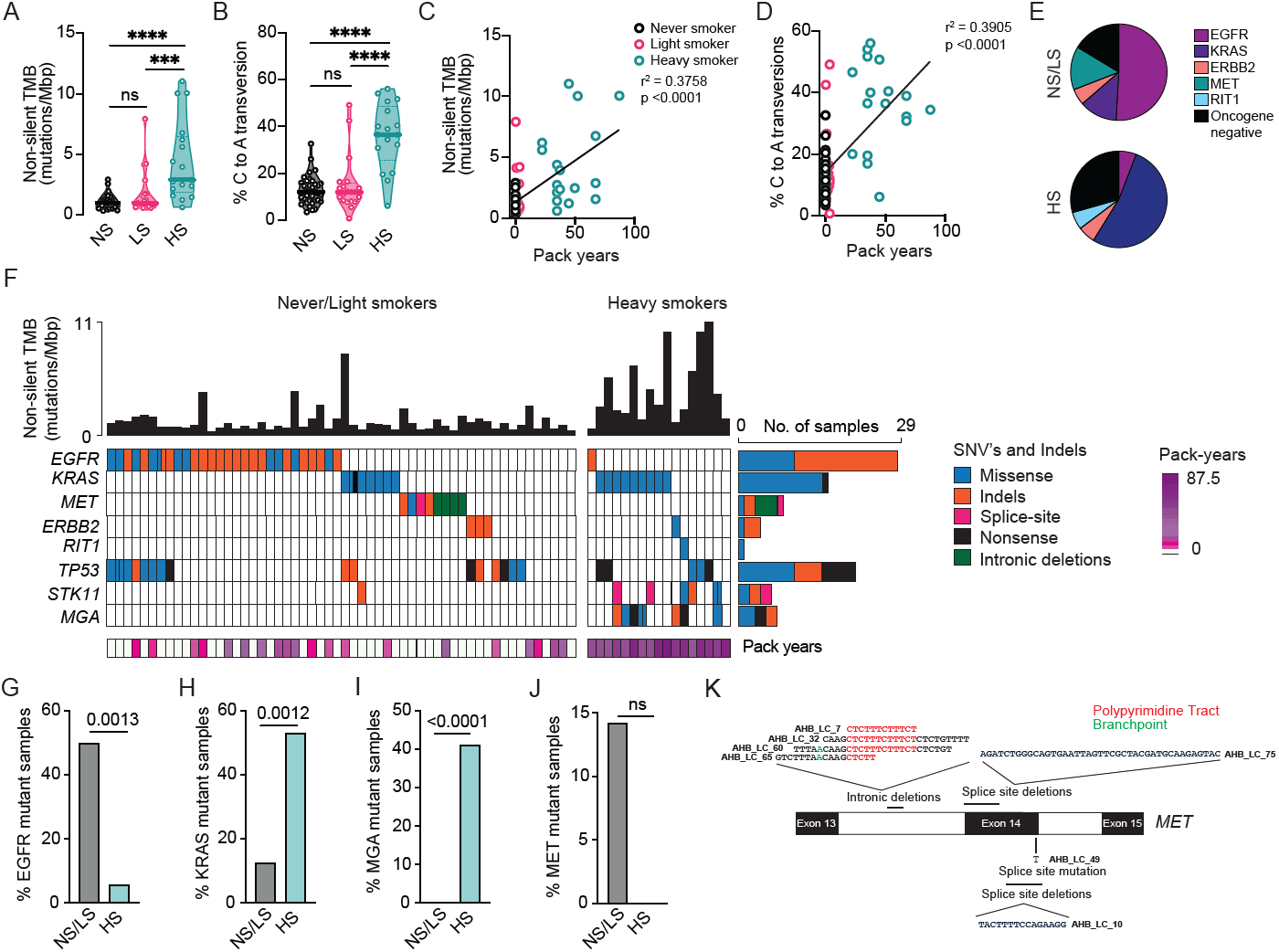
Altered somatic mutation rate and somatically mutated genes in never-smokers. **(A)** Non-silent tumor mutational burden (TMB) rate in never-smokers (NS) (< 100-lifetime cigarettes), light-smokers (LS) (<5 pack years), and heavy smokers (HS) (>20 pack years). Mann-Whitney test (two-tailed); **** p < 0.0001; ***p<0.001; ns p>0.05. **(B)** Percent of C to A transversions in NS, LS, and HS. Mann-Whitney test (two-tailed); **** p < 0.0001; ns p>0.05. **(C)** Association between non-silent TMB and pack-years smoked. Simple linear regression between non-silent TMB and pack-years; p < 0.0001. **(D)** Association between percent C to A transversions and quantitative pack years smoked. Simple linear regression; p < 0.0001. **(E)** Pie charts showing the number of oncogene-positive samples containing canonical driver mutations in the genes shown, and oncogene-negative samples in never-/light-smokers (NS/LS) and heavy smokers (HS). **(F)** Oncoplot of key genes belonging to the RTK/Ras/Raf pathway or genes known to be altered in lung adenocarcinoma. The figure shows known canonical mutations. The top bar plot shows the non-silent TMB rate for each patient. **(G-J)** The total number of samples with *EGFR, KRAS, MGA*, and *MET* mutations in NS/LS versus HS. Statistical analysis was done using two-tailed Fisher’s test. **(K)** Schematic representation of the *MET* locus between exon 13 and exon 15 and identified alterations likely to promote exon 14 skipping.

Mutations in the RTK-Ras-Raf pathway are critical drivers of lung adenocarcinoma^4^. We identified somatic mutations of genes in this pathway in 84% of tumors from never-/light-smokers and 71% of heavy smokers (**Figure 1E**). However, the proportion of samples with mutations in specific genes of the pathway varied between the groups (**Figure 1F**). *EGFR* mutations were more prevalent in never-/light-smokers (50% vs. 5.8%; Fisher’s exact test, p=0.0013) (**Figure 1G**) and *KRAS* mutations were enriched in heavy smokers (12.5% vs. 52.9%; Fisher’s exact test, p= 0.0012) (**Figure 1H**).

Mutations in the MYC transcription factor network member, *MGA*, were significantly associated with smoking history (**Figure 1I**). MGA regulates MYC-mediated transcription via its ability to dimerize with MAX and recruit a variant Polycomb complex^27^. Loss-of-function somatic mutations occur in ∼10% of human lung adenocarcinomas, and loss of *MGA* cooperates with mutant *KRAS* to promote lung cancer *in vivo*^*28*^. Interestingly, we found *MGA* mutations are exclusive to the tumors from smokers (41% of smokers), and significantly co-occur with *KRAS* mutations (4/7 or 57% of *MGA* mutations co-occur with mutant *KRAS*; Fisher’s exact test, p=0.0022; **Figure 1F**). Together these data identify tumor suppressor inactivation of *MGA* as a potential contributor to smoking-associated lung cancer in older women.

In recent years, somatic *MET* exon 14 skipping mutations have emerged as biomarkers for clinical response to *MET*-targeted therapies^4,29,30^. These variants disrupt the splice sites flanking exon 14, resulting in exon skipping and expression of a smaller isoform of *MET* with enhanced protein stability and kinase activity^31,32^. We observed *MET* exon skipping mutations at a higher prevalence than previous studies^4,29^, with a trend towards exclusivity in the never-/light-smoking group (8/56 14.2%; Fisher’s exact test, p=0.18) (**Figure 1J**). In addition to SNVs that disrupt canonical splice site motifs, intronic baits included in our exome panel allowed us to identify deletions in the upstream intron encompassing the intron 13-14 branchpoint or the polypyrimidine tract in four additional tumors^31,32^ (**Figure 1K**).

A possible explanation for the high prevalence of *MET* mutations in this cohort may be the advanced age of the participants (average age at diagnosis of 76-77 for never-/light-smokers) **(Table S3)**. Analysis of lung adenocarcinomas sequenced at Memorial Sloan-Kettering Cancer Center^33–36^ (MSKCC) showed that the age at diagnosis of patients with *MET*-mutant lung tumors was significantly higher compared to those with wild-type *MET* (mean age of *MET* mutant patients = 70.24 vs. Mean age of *MET* wild-type patients = 62.42, Mann Whitney test p=0.0002) (**Figure S1B**).

Several additional non-Ras pathway genes showed altered mutational burden in smokers and never-/light-smokers **(Figure S1C)**; these genes included *STK11*, also known as *LKB1*, which is well-known to have increased prevalence in smokers^37^ and confer a worse prognosis for tumors treated with PD-1/PD-L1 checkpoint inhibitors^38^. **(Figure 1F)**. *STK11* alterations occurred more frequently in heavy smokers than in never-/light-smokers (29% vs. 2%, Fisher’s exact test, p=0.0021). *STK11* alterations were predominantly mutually exclusive with *TP53* mutations (**Figure 1** and **Figure S1D**). We also noted the enrichment of *ATF7IP* somatic mutations in tumors from smokers (3/17 (18%) vs.0/56 (0%); Fisher’s exact test, p=0.01) (**Supplementary Figure S1E**). However, only three overall somatic mutations in *ATF7IP* were identified, and *ATF7IP* mutations have been previously observed in never-smokers^39^.

### Absence of lung cancer fusions events involving *RET, ROS1*, and *ALK*

Surprisingly, no canonical fusions involving *RET, ROS1*, or *ALK* were identified in any tumors. Typically, these fusions are prevalent in lung tumors from never-smokers. For example, we performed a meta-analysis of over 1000 patients with lung adenocarcinoma from ten genomic studies (**Methods**) and found that 15% of never-smokers have fusions involving *ALK, RET*, or *ROS1* compared to only 2.8% in moderate smokers (>5 pack years to ≤ 19 pack-years) and 1.16% in the heavy smoker (≥ 20 pack-years). Compared to historical data from MSKCC^33–36^, 14% of tumors would be expected to harbor fusions, versus the 0% observed in our cohort (**Supplementary Figure S2A**; p = 0.0028). To determine if alternative methods could identify fusions, we performed RNA amplicon sequencing on five oncogene-negative tumors and two controls **(Methods)**. While a *KIF5B-RET* and *EML4-ALK* fusion were readily detected in the positive controls, no fusions were detected in the five test tumors (**Supplementary Figure S2B**). We cannot exclude the possibility that fusions were missed for technical reasons. However, alternatively, the advanced age of this cohort might explain the absence of *RET, ROS1*, and *ALK* fusions, since kinase fusions have been associated with young age^40,41^. Notably, over 47% of the 112 participants in the Genomics of Young Lung Cancer study had tumors with *ALK, RET*, or *ROS1* fusions^15^ and our meta-analysis confirmed that fusions are more prevalent in tumors from younger individuals **(Supplementary Figure S2C)**. Moreover, mutational timing analysis identified chromosomal fusions as events that can occur early in life^42^, so fusions may be a general feature of lung tumors in younger patients rather than older individuals.

### Mutational processes of tumors in never-smokers include clock, APOBEC, and DDR deficiency

Mutational signatures in cancer can provide insight into cancer etiology and mechanisms of tumor therapy response^43^. Analysis of our cohort identified the presence of 17 known signatures from the COSMIC database (https://cancer.sanger.ac.uk/signatures/), accounting for a median of 90% of mutations in each sample (**Figure 2A and Table S5**). The predominant signatures were tobacco (SBS4), age-related clock-like process (SBS1 and SBS5), defective DNA damage response (SBS3, SBS6, SBS26, SBS30), and APOBEC mutagenesis (SBS2 and SBS13) (**Figure 2A**). Several signatures of unknown etiology were identified, but the contribution of these signatures to each mutational profile was low (about 3% of all somatic SNVs) (**Table S5**).

**Figure 2.**
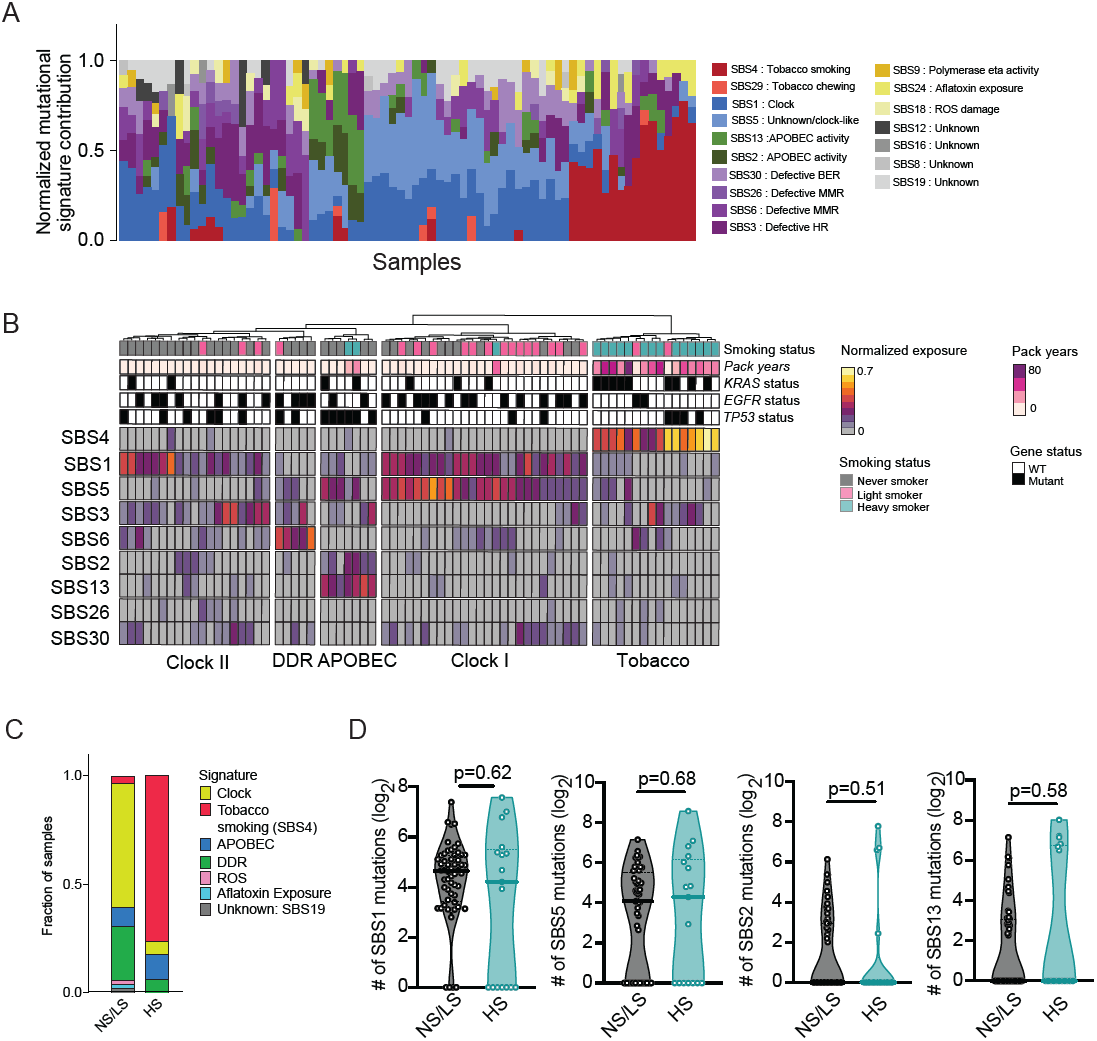
Somatic mutational signatures distinguish tumors from never-/light- and heavy smokers. **(A)** Contribution of each single base substitution (SBS) mutational signature to the total repertoire of mutational signatures identified in each patient. The fractional contribution is calculated by normalizing each signature exposure to the total signature exposure in each patient. Each stacked bar represents the mutational signature profile for an individual patient. **(B)** Heatmap of unsupervised clustering of normalized nine mutational signatures using Ward’s minimum variance method for both samples and signatures. The clustering is based on the normalized signature exposures. Features of each sample are shown above the heatmap including smoking status group, pack years, and mutational status of *EGFR, KRAS*, and *TP53*. **(C)** Bar graph indicating the mutational signature contributing to the maximal mutational burden for each sample. **(D)** Comparison of the number of mutations attributable to clock signatures (SBS1 and SBS5) and APOBEC signatures (SBS2 and SBS13) in never-/light-smokers and heavy smokers. Statistical significance was calculated using the Mann-Whitney test (two-tailed).

Unsupervised hierarchical clustering of mutational signature exposure identified five predominant mutational signature groups (**Figure 2B**). As expected, the tobacco/SBS4-high group was composed of 14/17 (82%) heavy smokers with SBS4 as the dominant mutagenic process accounting for an average of 41% of mutations in tumors of this group. Two (4%) light-smoker tumors also clustered into this group; one had an *EGFR* mutation and the other harbored mutant *KRAS*. The remaining 54 never- and light-smokers had little evidence of smoke exposure (SBS4 minimum and median fraction contribution = 0; SBS4 max fraction contribution 0.18), despite most individuals in the study reporting passive smoke exposure (**Figure S3A**). Thus, passive smoke exposure is not likely to be a major driver of mutagenesis in lung cancer in never-smokers.

Clock-like and DNA damage signatures dominated the mutagenic landscape of tumors in never-/light-smokers (**Figure 2A-B**). Clock-like signatures SBS1 and SBS5 were the predominant mutagenic process in 33 of 56 (59%) never-/light-smokers (**Figure 2C**). These clock-like signatures are believed to arise from mitotic errors, with the accumulation of clock-like mutations increasing with age^44^. We observed a significant correlation between SBS1 and SBS5 signatures (simple linear regression R-squared = 0.1208; p=0.0026), but a group of SBS5-low tumors was also evident, indicating that the two signatures may reflect related but distinct mutagenic processes (**Figure S3B**).

Seven tumors with elevated SBS2 and SBS13 signatures clustered in an APOBEC-high cluster. This group comprised five never-/light- and two heavy smokers, and was enriched for samples with mutant *TP53* (6/7; Fisher’s exact test, p=0.0018). The APOBEC mutational signature is characterized by C to T and C to G mutations due to elevated activity of the APOBEC enzymes with a polynucleotide cytosine deaminase activity^45–47^. APOBEC activity has been shown to be associated with the early onset of lung adenocarcinoma in east-Asian females never-smokers^48^.

To determine whether never-smokers had a higher overall burden (rather than proportion) of clock and APOBEC mutations, we estimated the absolute number of mutations attributable to each signature. This analysis revealed no significant difference in clock-like mutagenesis (SBS1 p = 0.62 and SBS5 p = 0.68 by Mann Whitney’s U-test) or APOBEC mutagenesis in tumors from never-/light- and heavy smokers (Mann Whitney SBS1 p = 0.51 and SBS5 p = 0.58) (**Figure 2D**). Therefore we conclude that these common mutagenic processes are operative in lung cells in general rather than disproportionately affecting smokers or never-smokers.

The last group of samples showed evidence of defective DNA damage repair (DDR), including tumors with either mismatch repair (MMR; SBS6) or homologous recombination (HR; SBS3) defect signatures. We sought to identify the drivers of these signatures by identifying somatic or germline variants belonging to known genes in these genome integrity pathways (**Methods**) but did not find any known mutations in these samples (**Figure S3C**). Analysis of germline variants did reveal four samples with heterozygous pathogenic germline mutations in *MUTYH* (**Figure S3D**). However, one of these samples had a dominant HR signature rather than the defective base excision repair signature expected from *MUTYH* deficiency. Further investigation is warranted to identify the underlying cause of DNA damage signatures in never-smokers and to determine if these mutations contribute to tumor initiation and therapeutic response.

### The mutation spectrum of *KRAS* and *EGFR* impacts therapeutic options in never- and light-smokers

Given the substantial differences in the mutational signatures between smokers and never-/light-smokers, we hypothesized that frequently mutated genes might also show differences in their mutational spectrums. Indeed, *EGFR* showed significant skewing of the type of mutations identified by smoking status; 63% of the *EGFR* mutations in the never-/light-smokers group were indel mutations in exon 19 or 20 rather than missense variants such as L858R^49^. Extending this analysis to the MSKCC^33–36^ and Campbell^50^ cohorts confirmed a higher proportion of *EGFR* indel mutations in never-/light-smokers compared to heavy smokers (**Figure 3A**). A possible explanation for an increased proportion of *EGFR* indels in never-/light-smokers could be altered DNA repair resulting in a genome-wide increase in insertion/deletion mutagenesis. However, all tumors in our cohort had a similar abundance of indel mutations regardless of their *EGFR* genotype (Mann Whitney test, indel vs. wild-type p=0.129, indel vs. missense p=0.5184) (**Figure S4A**), so indel mutagenesis *per se* may not be altered. Conversely, missense variants in *EGFR* could be due to smoking-related mutagenesis. However, the nucleotide changes resulting in the L858R variant are not characteristic of smoking-induced mutagenesis (**Figure S4B**), so the reason for this skewed mutational spectrum remains unclear.

**Figure 3.**
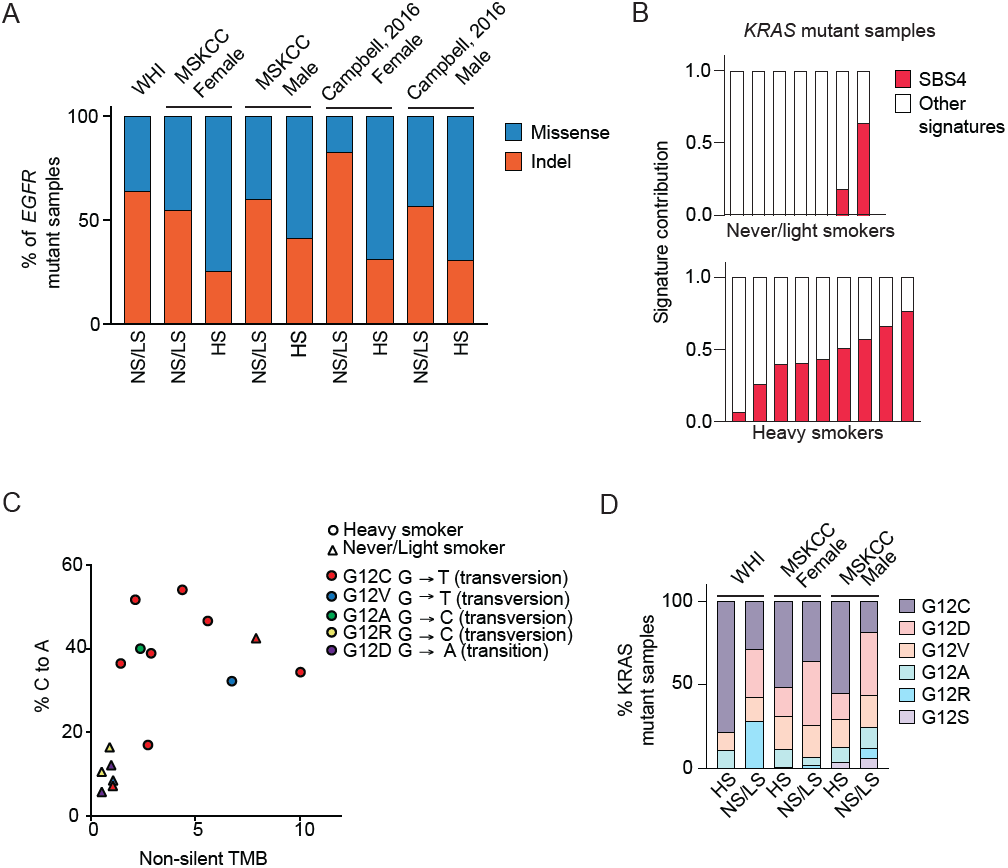
Enrichment of *EGFR* indel and specific *KRAS* variants in never-/light-smokers. **(A)** Percent indel and missense mutations in *EGFR* mutant samples in our cohort (WHI) and MSKCC^33–36^ and Campbell^50^ cohort cohorts. **(B)** SBS4/Tobacco smoke signature contribution to the total detectable mutational signature spectrum of samples with *KRAS* mutations. The contribution of SBS4 (red bars) is shown relative to the contribution of all other signatures (white bars). **(C)** Scatter plot demonstrating the association between non-silent tumor mutational burden and percent C to A transversions in *KRAS* mutant samples in never-/light-smokers (triangles) and heavy smokers (circles). **(D)** *KRAS* mutation spectrum at the amino acid level in the WHI and MSKCC cohort by the smoking group. Percentage of samples with specific *KRAS* G12 amino acid changes plotted as a percentage of all G12 alterations.

*KRAS* mutations are very frequent in tumors from smokers and to a lesser extent can be observed in tumors from never-smokers^21^. We identified relatively frequent mutation of *KRAS* in never-/light-smokers (n=7; 12.5%) in addition to the expected enrichment of *KRAS* mutations in heavy smokers (52.9%). To address if *KRAS* mutations occur in never-smokers due to secondary smoke exposure, we queried the levels of the SBS4 tobacco signature in the *KRAS*-mutant tumors. Heavy smokers with *KRAS* mutations uniformly exhibited an SBS4 tobacco smoking signature in their tumors (**Figure 3B, Table S5**). In contrast, the SBS4 signature exposure was below the detection level in five of seven tumors in the never-/light-smokers group (**Figure 3B**), so we conclude that *KRAS* mutations do occur in the absence of smoke exposure.

The predominant site for mutation in *KRAS* in lung cancer is glycine 12 and all *KRAS*-mutant tumors in our cohort were mutated at that site. However, the specific amino acid variant introduced differed between never-/light-smokers and heavy smokers (**Figure 3C**). *KRAS* mutations in tumors from smokers were predominantly G12C variants (7/9 or 78%), whereas never-/light-smokers had fewer G12C variants (2/7 or 28.5%) and a higher percentage of G12D variants both in our cohort as well as the MSKCC^33–36^ and Campbell^50^ studies (49% G12C mutations compared to only 22% in never-/light-smokers; Fisher’s exact test, p=0.0005 **Figure 3D; Figure S4C**). Currently, G12C is the only clinically druggable *KRAS* variant^51,52^, so these differences in genotype impact patients’ access to the newly available *KRAS-*targeted therapies.

### The genomic copy number landscape is similar between never and heavy smokers

Somatic copy number alterations (SCNAs) are a distinctive feature of lung cancer genomes^3,53–55^. SCNAs include focal amplifications and deletions, chromosome arm-level events, aneuploidy, and whole genome doubling^56,57^. Whereas the role of smoking on SNV mutagenesis is well documented, the impact of smoking on aneuploidy is not well understood. Ploidy or the number of complete sets of diploid genomes is frequently altered in cancer due to underlying errors in cell division^58^. These ploidy changes are often due to whole genome duplications, sometimes followed by secondary gains or losses of full or partial sets of chromosomes. Despite the extensive differences in tumor somatic mutation patterns between heavy and never-/light-smokers, we surprisingly found no significant difference in ploidy between these groups (Mann Whitney test, WHI cohort p=0.86; Campbell^50^ cohort: female p=0.23; male p=0.60; Fishers exact test p=0.13) (**Figure 4A, Figure S5A and Table S6**). Furthermore, ploidy did not correlate with pack-years of cigarettes smoked in both our cohort as well as the Campbell^50^ cohort (Spearman’s correlation WHI cohort: r=-0.06, p=0.612 and Campbell^50^ cohort: r=0.060, p=0.154) **(Figure S5B)**.

**Figure 4.**
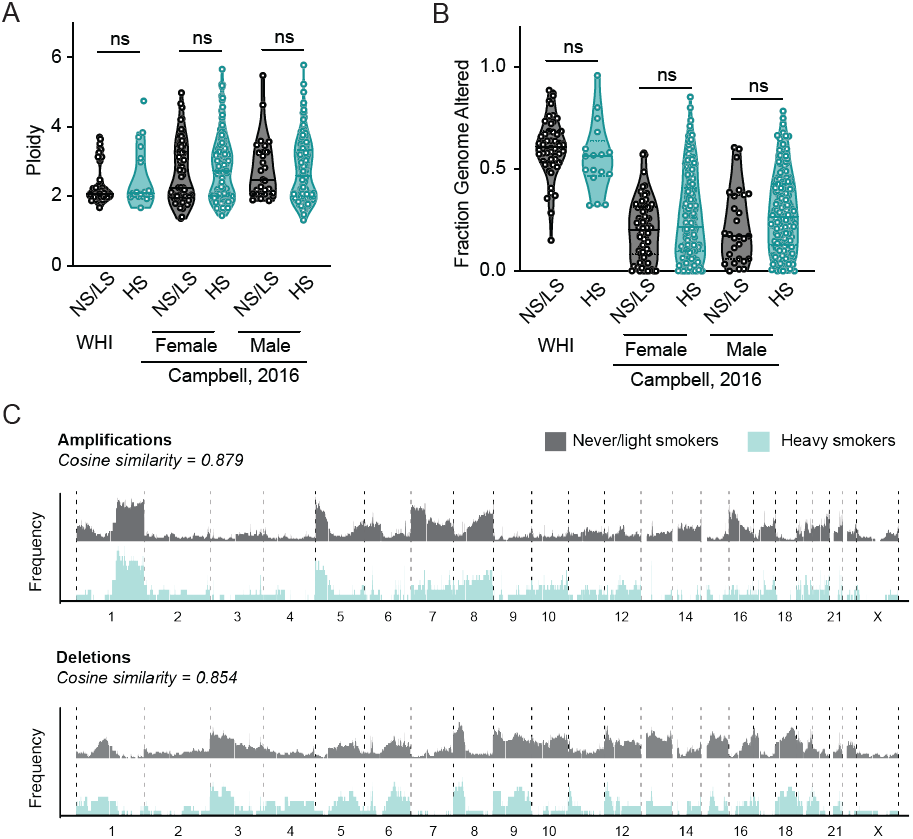
Somatic copy number changes do not differentiate tumors from never-/light-smokers and heavy smokers. **(A)** Genome ploidy of lung cancer patients in the WHI cohort and Campbell^50^ cohort study in never-/light-smokers (gray) and heavy smokers (blue/green). Mann-Whitney test (two-tailed); ns: not significant **(B)** Fraction genome altered (FGA) of never-/light-smokers and heavy smokers from the WHI and Campbell^50^ cohort cohorts. Mann-Whitney test (two-tailed); ns: not significant **(C)** Pattern of amplifications and deletions in never-/light-smokers (gray/top panels) and heavy smokers (blue/bottom panels) across all 23 chromosomes. Cosine similarity was calculated between both smoking groups for amplifications (top) and deletions (bottom).

We next examined the difference in the fraction genome altered (FGA) between never-/light-smokers and heavy smokers. Unlike ploidy, FGA describes the overall breadth of the genome altered rather than the amplitude of that alteration across the whole genome. There was no significant difference in the FGA between both never-/light- and heavy smokers in our cohort (p>0.05) (**Figure 4B, Methods)**. Analysis of the MSKCC^33–36^ cohort showed that tumors from never-/light-smokers paradoxically had higher FGA than heavy smokers (p<0.0001) **(Figure S5C**). Consistently, FGA did not correlate with pack-years of cigarettes smoked in our cohort (Spearman’s correlation r=-0.19, p=0.10) and showed a negative correlation in the Campbell cohort (Spearman’s correlation r=-0.08, p=0.02) **(Figure S5D)**. The total number of amplifications or deletions between both smoking groups was also not significantly different **(Figure S5E)**. We then compared the pattern of recurrent amplifications and deletions across the genome between never-/light- and heavy smokers (**Figure 4C, Methods**). Overall, the frequency of alterations at specific chromosomal locations showed broadly similar patterns. Therefore, smoking does not appear to influence the quantitative and qualitative metrics of genome-wide copy number patterns in lung adenocarcinoma.

### Arm-level copy number alterations cluster tumors independent of smoking status

SCNA burden is associated with poor overall survival and is being considered as a potential biomarker of recurrence and therapy^55,59–61^. Recent work showed that tumors from never-smokers contain frequent arm-level copy number alterations, and these can be used to cluster tumors into distinct groups with increasing aneuploidy^11^. Unsupervised clustering of arm-level copy number events **(Table S7 and Methods)** recapitulated three copy number groups similar to those previously described: Group I (n=14), Group II (n=38), and Group III (n=21) (**Figure 5A**). Group I tumors was enriched for deletions of most arm-level deletion events such as 3p, 9p, 17p which have been shown to be frequently lost across cancers^56^. Group II tumors included 52% (n=38) of all samples in the cohort with very few arm-level events. Group III showed significant enrichment of amplifications of 7p, 7q, 6p, and 20p compared to the other two subtypes combined. However, we did not observe amplification of 1q or 5p in this group, unlike the previous study^11^. Interestingly, smoking history did not appear to influence tumor clustering (**Figure 5B**). Consistently, there was no significant difference in non-silent tumor mutational burden between the three groups that would suggest a role for smoking in the tumor clustering (**Figure 5C**). We did observe the enrichment of *TP5*3-mutant samples in Group I (Fisher’s exact test p = 0.0183) and *KRAS*-mutant samples in Group II (Fisher’s exact test p = 0.0492). Whole genome duplication (or ploidy >2) was significantly higher in both Group I and III compared to Group II (Fisher’s exact test p<0.05) (**Figure 5D**), as well as FGA (Unpaired t-test p<0.05) (**Figure 5E**). Overall, we find that these aneuploidy-based clusters are independent of the patients smoking history, suggesting a non-smoking-related etiology.

**Figure 5.**
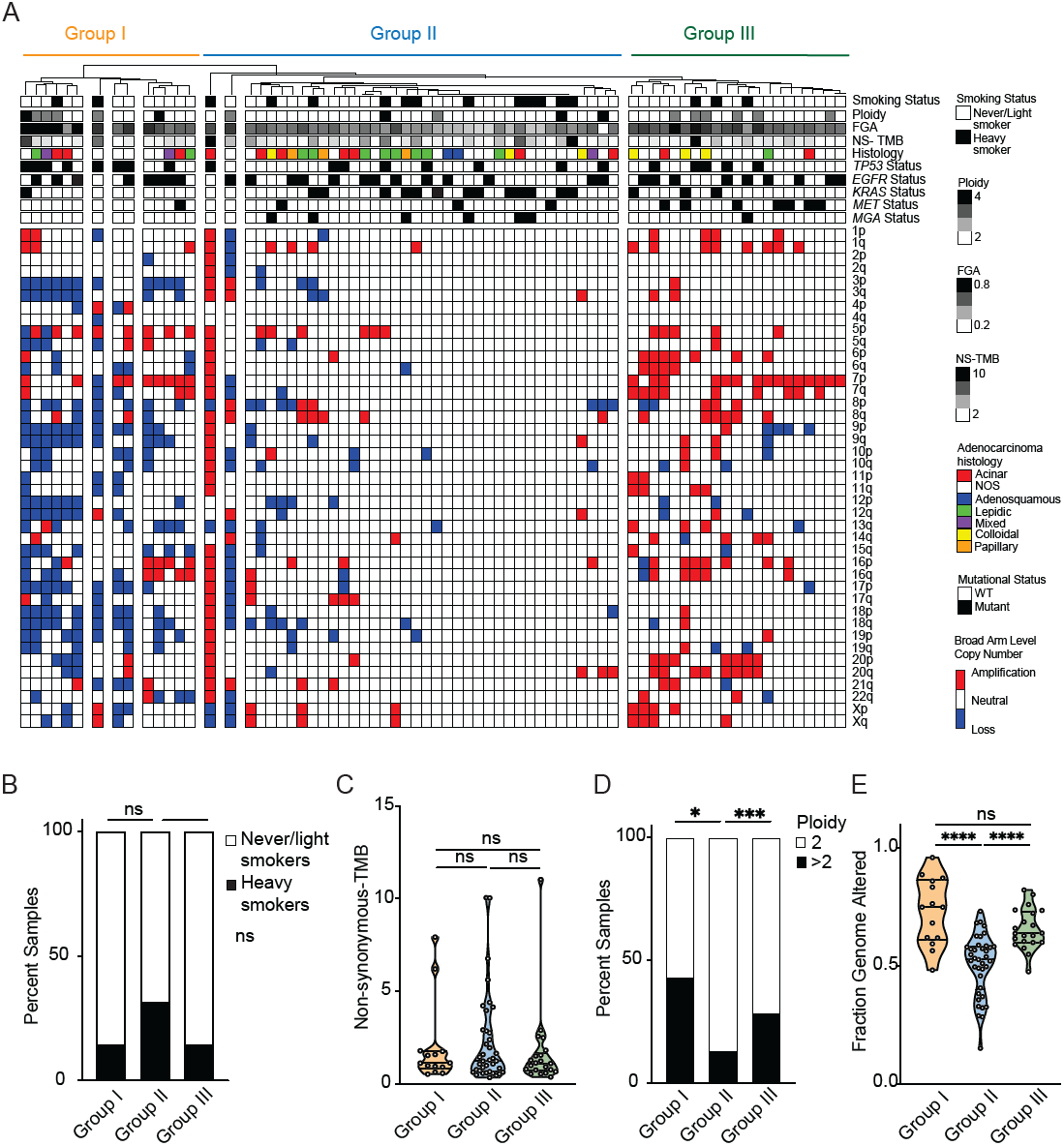
Arm-level copy number alterations cluster samples into groups unrelated to the history of smoking. **(A)** Heatmap of unsupervised clustering of arm-level copy number alterations in the WHI cohort using Ward’s minimum variance method for both samples and signatures. The clustering is based on binarized arm-level calls from GISTIC 2.0. Samples were grouped into three groups based on broad clusters and copy number patterns. **(B)** Stacked bar graph showing the percent of never-/smokers and heavy smokers in each copy-number group. **(C)** Non-silent TMB in samples split by arm-level copy number group. Mann Whitney test; ns: not significant. Group I: Orange; Group II: Blue and Group III: Green **(D)** Stacked bar graph showing percent samples in each group with ploidy 2 or ploidy greater than two. Black bars indicate a ploidy estimate greater than 2 and white bars indicate a ploidy estimate of 2. Fisher’s exact test. **(E)** Fraction genome altered in samples split by arm-level copy number group. Unpaired t-test, **** p<0.0001

## Discussion

In this study, we present the genomic landscape of lung adenocarcinoma in female never-/ light-smokers from the Women’s Health Initiative. Our work complements recent proteogenomic and genomic analyses^11,12,15,48,62^, but specifically focuses on older women, the most common demographic among never-smoker lung cancer cases. We confirm the well-known genomic differences between never-/light- and heavy smokers such as the enrichment of *EGFR* mutations in never-smokers. We also uncover unique features of the genomic landscape that have previously not been described, such as the enrichment of *MGA* mutations in heavy smokers. Our data also points to interesting age-related differences in the pathogenesis of lung cancer, with older never-smokers having a lower prevalence of RTK fusions and a higher prevalence of *MET* exon 14 skipping mutations. While somatic SNV and indel mutational patterns are distinct between never-/light-smokers and heavy smokers, surprisingly, we found no association between smoking status and aneuploidy/SCNAs. Therefore the recently described copy number subtypes of lung tumors in never-smokers^11^ are likely to represent a general segregation of tumors into copy number subtypes not unique to tumors from smokers or never-smokers.

Despite the similar patterns of aneuploidy, tumors from never-smokers and smokers show vast differences in somatic mutation landscape with important implications for therapy availability and efficacy. First, somatic tumor mutational burden, an indicator of neoantigen burden, is suggested to predict to a certain extent immunotherapy response^35,63^. Thus the low TMB of tumors from never-smokers may partially contribute to their immunologically “cold” phenotype^64^. Second, we show that the mutational spectrum of variants in important clinical targets *EGFR* and *KRAS* are significantly different in tumors from never-/light- and heavy smokers. *EGFR* indel mutations are enriched in tumors from never-/light-smokers while missense mutations are more common in heavy smokers. The reason for this difference is yet unclear, but it may be related to differences in DNA damage repair mechanisms or other mechanisms of mutagenesis and may be an important clue to the etiology of *EGFR*-mutant lung cancer. Interestingly, we also found *KRAS* mutations in tumors with no tobacco smoke exposure, but the targetable G12C mutation is much less prevalent in these tumors. Thus we suggest that never-/light-smokers with *KRAS*-mutant disease are a population with particular unmet medical needs; these tumors naturally lack other targetable biomarkers such as *EGFR* mutations, do not contain a targetable *KRAS* variant, and are unlikely to respond to immunotherapy. Fortunately, the development of additional *KRAS* inhibitors is underway to partially address this challenge.

Exome-wide mutational signatures provide a clear view of mutational processes at work, such as the distinctive SBS4 or “transversion-high” signature observed in tumors from smokers^17,50^. We reasoned that focused analysis of tumors from never-/light-smokers might reveal clues to mutagenic processes and the etiology of lung cancer in the absence of smoking. However, we observed no signatures that were uniquely abundant in tumors from never-/light-smokers. Age-related clock mutagenesis and APOBEC mutagenesis were clearly operative but contributed to a similar mutational burden in tumors from never-/light- and heavy smokers. Thus, we cannot attribute the cancer development in never-/light-smokers to any unique environmental or endogenous mutational process at this time. As the number of lung cancer cases in never-smokers appears to be increasing, it is crucial to continue to better understand the molecular mechanisms of tumor development in never-smokers to develop effective prevention and treatment strategies.

## Acknowledgments

Peggy Porter, Jamie Guenthoer, Meredith Bemus, Kyle Manning, Matthew Meyerson, John Minna, Navonil De Sarkar, Megan Skinner Herndon, Urvashi Pandey, Esther Jhingan, Sushma S. Thomas, Gretchen Van Lom, Adi Gazdar, Jian Carrot-Zhang, Robert Eisenman, James P. Adams, Jeffrey J. Delrow, Todd Panek, Stephanie J Weaver, Lisa G Johnson, Allison Matson, Elizabeth M. Donato, Michelle A. Wurscher, Lorna G. Nolan, Dana Farber Cancer Institute, Center for Cancer Genomics.

## Funding

This work was supported in part by the International Association for the Study of Lung Cancer and Lung Cancer Foundation of America; Prevent Cancer Foundation; Seattle Translational Tumor Research Lung Cancer Program; National Cancer Institute (NCI) R37CA252050 to A.H.B.; the Fred Hutch Translational Data Science Postdoctoral Fellowship to S.M.; the Genomics & Bioinformatics Shared Resource, <u>RRID:SCR_022606</u>, of the Fred Hutch/University of Washington Cancer Consortium (P30 CA015704); NCI K22CA237746 to G.H.

## Author contributions

Conceptualization: G.L.A., A.H.B.; Data acquisition: S.M., A.N, N.A.P, C.S., A.R.T, G.L.A, A.H.B; Methodology: S.M., A.P., M.K., A.C.H.H, A.N., N.A.P., F.W., M.P.F, C.S., A.R.T, G.H.; Analysis: S.M., A.P., F.W., M.P.F.,M.P; Writing – original draft: S.M., G.H., A.H.B; Writing – review & editing, all authors; Funding acquisition: S.M., A.H.B, G.L.A.

## Declaration of interests

The authors declare no competing interests.

## Methods

### Ethics declaration

All samples were obtained from patients after approval from the Fred Hutch Institutional Review Board (#8667) and appropriate informed consent from participants. Patient data including sample identifiers, patient identifiers, and metadata identifiers were de-identified from the authors. No identifiable private information was generated in this analysis.

### Patient selection

The participants of the current study were all post-menopausal women retrospectively selected from the WHI cohort. Initial selection criteria for participants included a lung adenocarcinoma diagnosis and smoking history of either less than 100 lifetime time cigarettes (never-smokers) or less than 5 pack-years (light-smokers) or greater than 20 pack-years (heavy smokers). All cohort participants were matched for cancer stage, diagnosis year, and tissue purity. Patient characteristics are provided in **Supplementary Table S1**.

### Pathology review and tissue samples

H&E slides from Formalin Fixed Paraffin Embedded (FFPE) tumors were generated for the participants fulfilling the above selection criteria. All tissue samples were generated from either diagnostic surgery/lobectomy/segmentectomy/resection procedure. These sections were reviewed by a pathologist for histological confirmation of the lung adenocarcinoma diagnosis and tumor content and purity (check for sufficiency for sequencing). 73 participants with sufficient tumor availability for sequencing were included in the present study (Supplementary Table S1 and S3). While the tumor source for this study was derived from FFPE-embedded tissue, each sample had a matched normal/control, derived from fresh frozen peripheral blood. To control for FFPE-induced change 10 tumor-adjacent normal samples from participants in the study were also included.

### Genomic DNA Isolation for sequencing

Tumor and tumor-adjacent normal FFPE tissues were macro-dissected guided by pathological review of sections. Genomic DNA was isolated using the QIAGEN QIAmp DNA FFPE kit (Cat. No, 56404) with some modifications. DNA from matched normal was derived from the buffy coats of prepared blood samples. A salting out method was used to purify the genomic DNA. Red blood cells were first lysed and washed out and then the white blood cell nuclei underwent lysis. Cellular proteins were precipitated and removed, followed by DNA precipitation.

### Custom Whole Exome Sequencing and pre-analysis processing

Custom whole exome sequencing was performed using DNA derived from tumor/normal FFPE tissue and fresh frozen peripheral blood. 250 ng of FFPE-derived DNA and 150 ng of fresh frozen blood DNA were used for library construction. Normalized genomic DNA was fragmented to an average size of 250 bp, and size-selected DNA was ligated to adapters. Libraries were pooled and sequenced to quantify library yields. Pooled libraries were then captured using the “POPv3.1_SV_ONLY” (Design ID 3191451) bait set ^65^. Hybrid captures were then sequenced on NovaSeq flow cells. Sequencing metrics are provided in Supplementary Table S2. Read pairs were aligned to the hg19 reference sequence using the Burrows-Wheeler Aligner^66^, and data were sorted and duplicate-marked using Picard tools. The alignments were further refined using the Genome Analysis Toolkit (GATK)^67,68^ for localized realignment around indel sites and recalibration of quality scores was also performed. The complete analysis pipeline for alignment can be found at https://github.com/FredHutch/tg-wdl-LILAC-workflow.

Tumor and matched normal DNA pairing were unknown prior to sequencing therefore a fingerprinting analysis was performed using 44 polymorphic loci to identify the pairing. Picard Tools GenotypeConcordance was used to calculate the concordance that a given test sample matches the sample being considered. This was performed on all pairwise combinations of samples in the cohort. The output of the pair-wise comparisons was then mapped to a concordance matrix, where concordance values above 4 standard deviations of the median concordance value for the cohort indicated a high likelihood that the samples match. Potential matches are manually reviewed and confirmed for accuracy from the WHI.

### Genomic data submission

The genomic data will be submitted under the parent WHI study # phs000200.

### Mutation calling

To define a high-confidence map of somatic single nucleotide variants (SNVs) and insertion/deletion variants (indels), we called mutations using a custom mutation calling strategy involving three somatic callers: MuTect2, Strelka, and SvABA. Analysis-ready BAM files were analyzed using GATK-MuTect 2 (version 4.1.4.0) run with the FFPE bias filter and a Panel of Normals (PoN) including 10 FFPE normal samples to help exclude potential FFPE artifacts^69^. BAM files were also processed through Strelka (enter version)^70^ with Manta (enter version) ^71^. And BAMs were also evaluated using the SvABA algorithm^72^. The complete analysis pipeline for alignment and somatic SNV and indel calling using MuTect2 and Strelka can be found at https://github.com/FredHutch/tg-wdl-LILAC-workflow. SNV calls that passed both MuTect2 and Strelka were included in the final call set. Indel calls that passed at least two of the three callers were included in the final call set (Supplementary Table S4). Finally, those SNVs and indels with variant allele frequencies (VAF) greater than 10% in gnomAD or ExAC databases were filtered out to generate the final call set for further analysis. Significant SNV and indel mutations were identified using the MutSig2CV^69,73^ algorithm.

### Fusion detection and mutational analysis from targeted re-sequencing by AmpliSeq

To perform deeper targeted sequencing, we utilized Illumina’s AmpliSeq Focus Panel to identify missed mutational calls in otherwise oncogene-negative samples and fusions. For the DNA sequencing panel, genomic DNA extracted for custom whole exome sequencing was used as input for the targeted sequencing. RNA was extracted from FFPE slides/curls using the QIAGEN RNeasy FFPE Kit (Cat No. 73504).

All amplicon samples were sequenced on an Illumina MiSeq using a paired-end 150bp read configuration. Raw data were collected using Illumina Real Time Analysis (RTA) software 1.18.54.4, with subsequent base calling and demultiplexing performed with bcl2fastq 2.20 (https://support.illumina.com/sequencing/sequencing_software/bcl2fastq-conversion-software.html). All samples were sequenced to an average depth of 300K read pairs each.

For DNA amplicon analysis, AmpliSeq samples were processed with the Illumina DNA Amplicon Workflow v3.0.0.14, which internally uses BWA MEM 0.7.9a^66^to align paired reads to the GRCh37/hg19 human reference, followed by variant calling with Pisces in targeted amplicon region. Resulting DNA variants were annotated with GATK (v4.1.8.1) Funcotator along with funcotator_dataSources.v1.6.20190124s^68^.

For RNA fusion analysis, AmpliSeq samples were processed with the Illumina RNA Amplicon Workflow 3.0.0.26. This analysis method also uses BWA MEM internally to align reads to targeted regions and fusions, followed by proprietary methods to call gene fusions and exon variants.

### Analysis of external datasets

External genomic datasets were used to validate the findings made in the present study. All data were derived from cbioportal and no secondary analysis was performed. All data referred to as Campbell cohort was derived from^50^. MSKCC^33–36^ data was derived from cbioportal lung adenocarcinoma studies^33–36^. For fusion analysis data from MSKCC ^33–36^ OncoSG^74^, TCGA (TCGA Research Network: https://www.cancer.gov/tcga), Broad^3^ and Sherlock^11^ data was utilized.

### Mutational signature analysis

Mutational signature analysis was performed using the final variant call file generated for SNVs and indels. The R-based package Sigminer utilizes a non-negaive matrix factorization (NMF) based approach for mutational signature determination^43,75–85^. First, a manual *de novo* signature extraction was performed and then matched to known COSMIC signatures. Next, the same variant call file was used to perform an automatic signature extraction followed by matching to known COSMIC v3.1 signatures. Both signature calls were compared for their exposure and those signatures that showed an exposure of greater than 0.6 in any patient were used to generate a list of signatures of significance. The signatures of significance were then used to perform a final signature extraction limited to these signatures alone to form the final mutational signature exposure matrix. Unsupervised clustering was then performed of these mutational signature exposures using the Ward.D.2 minimum variance method using Euclidean distance.

### Copy Number Analysis

To determine the copy number alterations, we utilized the TITAN pipeline (https://github.com/gavinha/TitanCNA; commit 2b55d94)^86,87^. Corrected read counts were determined in non-overlapping windows of 50_kb that also overlapped the bait intervals by at least 1 bp. We also utilized the 10 FFPE “normal” samples to normalize any FFPE-induced copy number changes. All TITAN calls were then subject to manual curation to verify the optimal ploidy solution. Curated optimal solutions are shown in Table S6. Copy number data from TITAN was then input into GISTIC 2.0 to determine regions with significant copy number alterations. Fraction genome altered was calculated by dividing the sum of all amplified/deleted segments by the total number of segments for each patient. Arm-level copy number was determined from GISTIC output and unsupervised clustering was performed using Ward’s minimum variance method.

## Supplementary figure legends

**(S1A)** Number of female never-smokers diagnosed with lung cancer stratified by age at diagnosis from Siegel *et al*., 2021. **(S1B)** Age at diagnosis of *MET* wild-type and *MET* mutant samples from the MSKCC^33–36^ cohort. Median values for both groups are shown in red and quartiles are shown as black lines. Mann Whitney test ***p=0.0002 **(S1C)** Oncoplot of other genes belonging to the RTK/Ras/Raf pathway or genes known to be altered in lung adenocarcinoma. The figure shows both canonical and non-canonical mutations. The top bar plot shows the non-silent TMB rate for each patient. **(S1D)** Oncoplot of *TP53* and *STK11* mutations in the WHI cohort. **(S1E)** Mirror plot showing the percent samples with mutations in a few key genes. Genes that were significantly enriched in one or the other smoking groups were marked by a green star. Fisher’s exact test was used to measure enrichment.

**(S2A)** Pie chart showing expected and observed proportions of *ALK, RET*, and *ROS1* translocations. Expected observations were based on previously reported proportions in patients and the observed proportions were based on the WHI cohort; p=0.0028. **(S2B)** Schematic representation of translocations found in control cell line *H2228* and lung adenocarcinoma metastatic PDX (AHB001) sample. Schematic shows the location of regions involved in the translocation on the origin chromosomes, coverage of different regions of the genes involved, and the final structure of the translocation. **(S2C)** Percent of lung cancer cases diagnosed with *ALK, RET*, and *ROS1* translocations grouped by age at diagnosis. Open circles show the percent of patients diagnosed in a particular age range with no mutations in *ALK, RET*, or *ROS1* (colored by gene) while filled circles show the percent of patients with a translocation.

**(S3A)** Heatmap showing primary smoke exposure and second-hand smoke exposure (SHSe) for each patient. Samples are sorted based on increasing pack years. Also shown is the signature with the maximum contribution to the mutational signature burden. **(S3B)** Scatter plot for SBS1 versus SBS5 contributions per sample. Simple linear regression shows a significant association between SBS1 and SBS5 contributions in the WHI cohort; p = 0.0026. **(S3C)** Oncoplot for somatic and germline mutations in DNA damage and repair (DDR) genes. Samples are grouped based on clustering shown in **Figure 2B. (S3D)** Relative mutational profile in all possible 5’ and 3’ contexts in *MUTYH* mutated samples. Y-axis shows the relative contributions of each mutational context (X-axis).

**(S4A)** The number of indel mutations in *EGFR* indel mutant, *EGFR* missense mutant, and *EGFR* wild type in the WHI cohort. Mann Whitney test, ns: not significant **(S4B)** Schematic of *EGFR* L858R mutation and *KRAS* G12C mutation at the nucleotide level (top). Mutational spectrum of *EGFR* and *KRAS* mutant samples in the WHI cohort with either L858R or G12C mutations only (bottom). **(S4C)** *KRAS* mutation spectrum at the amino acid level in the Campbell^50^ cohort by the smoking group. Percentage of samples with specific *KRAS* G12 amino acid changes plotted as a percentage of all G12 alterations found in samples.

**(S5A)** Stacked bar graphs showing the percentage of samples with whole genome duplication (WGD) in never-/light-smokers (NS/LS) and heavy smokers (HS) from the MSKCC^33–36^ cohort. Enrichment of WGD in NS/LS or HS was measured with a two-tailed Fishers test. p = 0.1318 (not significant) **(S5B)** Scatterplot of ploidy versus pack-years cigarettes smoked in both the WHI cohort as well as the Campbell^50^ cohort. No significant trends were observed on applying simple linear regression. **(S5C)** Fraction genome altered in never-/light-smokers and heavy smokers in female and male patients with lung cancer from the MSKCC^33–36^ cohort. Mann-Whitney test (two-tailed); **** <0.0001. **(S5D)** Scatterplot of ploidy versus pack-years cigarettes smoked in both the WHI cohort as well as the Campbell^50^ cohort. No significant trend was observed on applying simple linear regression. **(S5E)** The total number of amplifications and deletions in the WHI cohort in never-/light-smokers and heavy smokers. Mann-Whitney test (two-tailed); ns: not significant

## Notes

### Competing Interest Statement

The authors have declared no competing interest.

## References

1. Sung H, Ferlay J, Siegel RL, Laversanne M, Soerjomataram I, Jemal A, et al. Global Cancer Statistics 2020: GLOBOCAN Estimates of Incidence and Mortality Worldwide for 36 Cancers in 185 Countries. CA Cancer J Clin. 2021 May;71(3):209–49.

2. Melosky B, Wheatley-Price P, Juergens RA, Sacher A, Leighl NB, Tsao MS, et al. The rapidly evolving landscape of novel targeted therapies in advanced non-small cell lung cancer. Lung Cancer. 2021 Oct;160:136–51.

3. Imielinski M, Berger AH, Hammerman PS, Hernandez B, Pugh TJ, Hodis E, et al. Mapping the hallmarks of lung adenocarcinoma with massively parallel sequencing. Cell. 2012 Sep 14;150(6):1107–20.

4. Cancer Genome Atlas Research Network. Comprehensive molecular profiling of lung adenocarcinoma. Nature. 2014 Jul 31;511(7511):543–50.

5. Sun S, Schiller JH, Gazdar AF. Lung cancer in never smokers--a different disease. Nat Rev Cancer. 2007 Oct;7(10):778–90.

6. Wakelee HA, Chang ET, Gomez SL, Keegan TH, Feskanich D, Clarke CA, et al. Lung cancer incidence in never smokers. J Clin Oncol. 2007 Feb 10;25(5):472–8.

7. Youlden DR, Cramb SM, Baade PD. The International Epidemiology of Lung Cancer: geographical distribution and secular trends. J Thorac Oncol. 2008 Aug;3(8):819–31.

8. Pelosof L, Ahn C, Gao A, Horn L, Madrigales A, Cox J, et al. Proportion of Never-Smoker Non-Small Cell Lung Cancer Patients at Three Diverse Institutions. J Natl Cancer Inst [Internet]. 2017 Jan;109(7). Available from: http://dx.doi.org/10.1093/jnci/djw295

9. Jemal A, Miller KD, Ma J, Siegel RL, Fedewa SA, Islami F, et al. Higher Lung Cancer Incidence in Young Women Than Young Men in the United States. N Engl J Med. 2018 May 24;378(21):1999–2009.

10. Rudin CM, Avila-Tang E, Samet JM. Lung cancer in never smokers: a call to action. Clin Cancer Res. 2009 Sep 15;15(18):5622–5.

11. Zhang T, Joubert P, Ansari-Pour N, Zhao W, Hoang PH, Lokanga R, et al. Genomic and evolutionary classification of lung cancer in never smokers. Nat Genet. 2021 Sep;53(9):1348–59.

12. Govindan R, Ding L, Griffith M, Subramanian J, Dees ND, Kanchi KL, et al. Genomic landscape of non-small cell lung cancer in smokers and never-smokers. Cell. 2012 Sep 14;150(6):1121–34.

13. Domagala-Kulawik J, Trojnar A. Lung cancer in women in 21th century. J Thorac Dis. 2020 Aug;12(8):4398–410.

14. Ragavan MV, Patel MI. Understanding sex disparities in lung cancer incidence: are women more at risk? Lung Cancer Manag. 2020 Jun 22;9(3):LMT34.

15. Gitlitz BJ, Novello S, Vavalà T, Bittoni M, Sable-Hunt A, Pavlick D, et al. The genomics of young lung cancer: Comprehensive tissue genomic analysis in patients under 40 with lung cancer. JTO Clinical and Research Reports. 2021 Jul;2(7):100194.

16. Siegel DA, Fedewa SA, Henley SJ, Pollack LA, Jemal A. Proportion of Never Smokers Among Men and Women With Lung Cancer in 7 US States. JAMA Oncol. 2021 Feb 1;7(2):302–4.

17. Alexandrov LB, Ju YS, Haase K, Van Loo P, Martincorena I, Nik-Zainal S, et al. Mutational signatures associated with tobacco smoking in human cancer. Science. 2016 Nov 4;354(6312):618–22.

18. Daoud A, Chu QS. Targeting Novel but Less Common Driver Mutations and Chromosomal Translocations in Advanced Non-Small Cell Lung Cancer. Front Oncol. 2017 Sep 29;7:222.

19. Pan Y, Zhang Y, Li Y, Hu H, Wang L, Li H, et al. ALK, ROS1 and RET fusions in 1139 lung adenocarcinomas: a comprehensive study of common and fusion pattern-specific clinicopathologic, histologic and cytologic features. Lung Cancer. 2014 May;84(2):121–6.

20. Russo A, Lopes AR, Scilla K, Mehra R, Adamo V, Oliveira J, et al. NTRK and NRG1 gene fusions in advanced non-small cell lung cancer (NSCLC). Precis Canc Med. 2020 Jun;3:14–14.

21. Riely GJ, Kris MG, Rosenbaum D, Marks J, Li A, Chitale DA, et al. Frequency and distinctive spectrum of KRAS mutations in never smokers with lung adenocarcinoma. Clin Cancer Res. 2008 Sep 15;14(18):5731–4.

22. Anderson GL, Manson J, Wallace R, Lund B, Hall D, Davis S, et al. Implementation of the Women’s Health Initiative study design. Ann Epidemiol. 2003 Oct;13(9 Suppl):S5–17.

23. Van Herck Y, Feyaerts A, Alibhai S, Papamichael D, Decoster L, Lambrechts Y, et al. Is cancer biology different in older patients? Lancet Healthy Longev. 2021 Oct;2(10):e663–77.

24. Sacher AG, Dahlberg SE, Heng J, Mach S, Jänne PA, Oxnard GR. Association Between Younger Age and Targetable Genomic Alterations and Prognosis in Non-Small-Cell Lung Cancer. JAMA Oncol. 2016 Mar;2(3):313–20.

25. Design of the Women’s Health Initiative clinical trial and observational study. The Women’s Health Initiative Study Group. Control Clin Trials. 1998 Feb;19(1):61–109.

26. Paskett ED, Caan BJ, Johnson L, Bernardo BM, Young GS, Pennell ML, et al. The Women’s Health Initiative (WHI) Life and Longevity After Cancer (LILAC) Study: Description and Baseline Characteristics of Participants. Cancer Epidemiol Biomarkers Prev. 2018 Feb;27(2):125–37.

27. Carroll PA, Freie BW, Mathsyaraja H, Eisenman RN. The MYC transcription factor network: balancing metabolism, proliferation and oncogenesis. Front Med. 2018 Aug;12(4):412–25.

28. Mathsyaraja H, Catchpole J, Freie B, Eastwood E, Babaeva E, Geuenich M, et al. Loss of MGA repression mediated by an atypical polycomb complex promotes tumor progression and invasiveness. Elife [Internet]. 2021 Jul 8;10. Available from: http://dx.doi.org/10.7554/eLife.64212

29. Frampton GM, Ali SM, Rosenzweig M, Chmielecki J, Lu X, Bauer TM, et al. Activation of MET via diverse exon 14 splicing alterations occurs in multiple tumor types and confers clinical sensitivity to MET inhibitors. Cancer Discov. 2015 Aug;5(8):850–9.

30. Lu X, Peled N, Greer J, Wu W, Choi P, Berger AH, et al. MET Exon 14 Mutation Encodes an Actionable Therapeutic Target in Lung Adenocarcinoma. Cancer Res. 2017 Aug 15;77(16):4498–505.

31. Kong-Beltran M, Seshagiri S, Zha J, Zhu W, Bhawe K, Mendoza N, et al. Somatic mutations lead to an oncogenic deletion of met in lung cancer. Cancer Res. 2006 Jan 1;66(1):283–9.

32. Onozato R, Kosaka T, Kuwano H, Sekido Y, Yatabe Y, Mitsudomi T. Activation of MET by gene amplification or by splice mutations deleting the juxtamembrane domain in primary resected lung cancers. J Thorac Oncol. 2009 Jan;4(1):5–11.

33. Martínez-Pérez E, Molina-Vila MA, Marino-Buslje C. Panels and models for accurate prediction of tumor mutation burden in tumor samples. NPJ Precis Oncol. 2021 Apr 13;5(1):31.

34. Jordan EJ, Kim HR, Arcila ME, Barron D, Chakravarty D, Gao J, et al. Prospective Comprehensive Molecular Characterization of Lung Adenocarcinomas for Efficient Patient Matching to Approved and Emerging Therapies. Cancer Discov. 2017 Jun;7(6):596–609.

35. Rizvi NA, Hellmann MD, Snyder A, Kvistborg P, Makarov V, Havel JJ, et al. Cancer immunology. Mutational landscape determines sensitivity to PD-1 blockade in non-small cell lung cancer. Science. 2015 Apr 3;348(6230):124–8.

36. Caso R, Sanchez-Vega F, Tan KS, Mastrogiacomo B, Zhou J, Jones GD, et al. The Underlying Tumor Genomics of Predominant Histologic Subtypes in Lung Adenocarcinoma. J Thorac Oncol. 2020 Dec;15(12):1844–56.

37. Pécuchet N, Laurent-Puig P, Mansuet-Lupo A, Legras A, Alifano M, Pallier K, et al. Different prognostic impact of STK11 mutations in non-squamous non-small-cell lung cancer. Oncotarget. 2017 Apr 4;8(14):23831–40.

38. Skoulidis F, Goldberg ME, Greenawalt DM, Hellmann MD, Awad MM, Gainor JF, et al. STK11/LKB1 Mutations and PD-1 Inhibitor Resistance in KRAS-Mutant Lung Adenocarcinoma. Cancer Discov. 2018 Jul;8(7):822–35.

39. Pros E, Saigi M, Alameda D, Gomez-Mariano G, Martinez-Delgado B, Alburquerque-Bejar JJ, et al. Genome-wide profiling of non-smoking-related lung cancer cells reveals common RB1 rearrangements associated with histopathologic transformation in EGFR-mutant tumors. Ann Oncol. 2020 Feb;31(2):274–82.

40. Takeuchi K, Soda M, Togashi Y, Suzuki R, Sakata S, Hatano S, et al. RET, ROS1 and ALK fusions in lung cancer. Nat Med. 2012 Feb 12;18(3):378–81.

41. Shaw AT, Hsu PP, Awad MM, Engelman JA. Tyrosine kinase gene rearrangements in epithelial malignancies. Nat Rev Cancer. 2013 Nov;13(11):772–87.

42. Lee JJK, Park S, Park H, Kim S, Lee J, Lee J, et al. Tracing Oncogene Rearrangements in the Mutational History of Lung Adenocarcinoma. Cell. 2019 Jun 13;177(7):1842–57.e21.

43. Alexandrov LB, Kim J, Haradhvala NJ, Huang MN, Tian Ng AW, Wu Y, et al. The repertoire of mutational signatures in human cancer. Nature. 2020 Feb;578(7793):94–101.

44. Alexandrov LB, Jones PH, Wedge DC, Sale JE, Campbell PJ, Nik-Zainal S, et al. Clock-like mutational processes in human somatic cells. Nat Genet. 2015 Dec;47(12):1402–7.

45. Alexandrov LB, Nik-Zainal S, Wedge DC, Aparicio SAJR, Behjati S, Biankin AV, et al. Signatures of mutational processes in human cancer. Nature. 2013 Aug 22;500(7463):415–21.

46. Cho RJ, Alexandrov LB, den Breems NY, Atanasova VS, Farshchian M, Purdom E, et al. APOBEC mutation drives early-onset squamous cell carcinomas in recessive dystrophic epidermolysis bullosa. Sci Transl Med [Internet]. 2018 Aug 22;10(455). Available from: http://dx.doi.org/10.1126/scitranslmed.aas9668

47. Roberts SA, Lawrence MS, Klimczak LJ, Grimm SA, Fargo D, Stojanov P, et al. An APOBEC cytidine deaminase mutagenesis pattern is widespread in human cancers. Nat Genet. 2013 Sep;45(9):970–6.

48. Chen YJ, Roumeliotis TI, Chang YH, Chen CT, Han CL, Lin MH, et al. Proteogenomics of Non-smoking Lung Cancer in East Asia Delineates Molecular Signatures of Pathogenesis and Progression. Cell. 2020 Jul 9;182(1):226–44.e17.

49. Robichaux JP, L. X, Vijayan RSK, Hicks JK, Heeke S, Elamin YY, et al. Structure-based classification predicts drug response in EGFR-mutant NSCLC. Nature. 2021 Sep;597(7878):732–7.

50. Campbell JD, Alexandrov A, Kim J, Wala J, Berger AH, Pedamallu CS, et al. Distinct patterns of somatic genome alterations in lung adenocarcinomas and squamous cell carcinomas. Nat Genet. 2016 Jun;48(6):607–16.

51. Hallin J, Engstrom LD, Hargis L, Calinisan A, Aranda R, Briere DM, et al. The KRASG12C Inhibitor MRTX849 Provides Insight toward Therapeutic Susceptibility of KRAS-Mutant Cancers in Mouse Models and Patients. Cancer Discov. 2020 Jan;10(1):54–71.

52. Canon J, Rex K, Saiki AY, Mohr C, Cooke K, Bagal D, et al. The clinical KRAS(G12C) inhibitor AMG 510 drives anti-tumour immunity. Nature. 2019 Nov;575(7781):217–23.

53. Weir BA, Woo MS, Getz G, Perner S, Ding L, Beroukhim R, et al. Characterizing the cancer genome in lung adenocarcinoma. Nature. 2007 Dec 6;450(7171):893–8.

54. Holland AJ, Cleveland DW. Boveri revisited: chromosomal instability, aneuploidy and tumorigenesis. Nat Rev Mol Cell Biol. 2009 Jul;10(7):478–87.

55. Kou F, Wu L, Zhu Y, Li B, Huang Z, Ren X, et al. Somatic copy number alteration predicts clinical benefit of lung adenocarcinoma patients treated with cytokine-induced killer plus chemotherapy. Cancer Gene Ther. 2022 Aug;29(8-9):1153–9.

56. Beroukhim R, Mermel CH, Porter D, Wei G, Raychaudhuri S, Donovan J, et al. The landscape of somatic copy-number alteration across human cancers. Nature. 2010 Feb 18;463(7283):899– 905.

57. Harbers L, Agostini F, Nicos M, Poddighe D, Bienko M, Crosetto N. Somatic Copy Number Alterations in Human Cancers: An Analysis of Publicly Available Data From The Cancer Genome Atlas. Front Oncol. 2021 Jul 28;11:700568.

58. Bielski CM, Zehir A, Penson AV, Donoghue MTA, Chatila W, Armenia J, et al. Genome doubling shapes the evolution and prognosis of advanced cancers. Nat Genet. 2018 Aug;50(8):1189–95.

59. Han X, Tan Q, Yang S, Li J, Xu J, Hao X, et al. Comprehensive Profiling of Gene Copy Number Alterations Predicts Patient Prognosis in Resected Stages I-III Lung Adenocarcinoma. Front Oncol. 2019 Aug 6;9:556.

60. Davoli T, Uno H, Wooten EC, Elledge SJ. Tumor aneuploidy correlates with markers of immune evasion and with reduced response to immunotherapy. Science [Internet]. 2017 Jan 20;355(6322). Available from: http://dx.doi.org/10.1126/science.aaf8399

61. Danielsen HE, Pradhan M, Novelli M. Revisiting tumour aneuploidy - the place of ploidy assessment in the molecular era. Nat Rev Clin Oncol. 2016 May;13(5):291–304.

62. Devarakonda S, Li Y, Martins Rodrigues F, Sankararaman S, Kadara H, Goparaju C, et al. Genomic Profiling of Lung Adenocarcinoma in Never-Smokers. J Clin Oncol. 2021 Nov 20;39(33):3747–58.

63. Rizvi H, Sanchez-Vega F, La K, Chatila W, Jonsson P, Halpenny D, et al. Molecular Determinants of Response to Anti–Programmed Cell Death (PD)-1 and Anti–Programmed Death-Ligand 1 (PD-L1) Blockade in Patients With Non–Small-Cell Lung Cancer Profiled With Targeted Next-Generation Sequencing. J Clin Orthod. 2018 Mar 1;36(7):633–41.

64. Dai L, Jin B, Liu T, Chen J, Li G, Dang J. The effect of smoking status on efficacy of immune checkpoint inhibitors in metastatic non-small cell lung cancer: A systematic review and meta-analysis. EClinicalMedicine. 2021 Aug;38:100990.

65. Hanna GJ, Lizotte P, Cavanaugh M, Kuo FC, Shivdasani P, Frieden A, et al. Frameshift events predict anti-PD-1/L1 response in head and neck cancer. JCI Insight [Internet]. 2018 Feb 22;3(4). Available from: http://dx.doi.org/10.1172/jci.insight.98811

66. Li H, Durbin R. Fast and accurate short read alignment with Burrows-Wheeler transform. Bioinformatics. 2009 Jul 15;25(14):1754–60.

67. McKenna A, Hanna M, Banks E, Sivachenko A, Cibulskis K, Kernytsky A, et al. The Genome Analysis Toolkit: a MapReduce framework for analyzing next-generation DNA sequencing data. Genome Res. 2010 Sep;20(9):1297–303.

68. DePristo MA, Banks E, Poplin R, Garimella KV, Maguire JR, Hartl C, et al. A framework for variation discovery and genotyping using next-generation DNA sequencing data. Nat Genet. 2011 May;43(5):491–8.

69. Benjamin D, Sato T, Cibulskis K, Getz G, Stewart C, Lichtenstein L. Calling Somatic SNVs and Indels with Mutect2 [Internet]. bioRxiv. 2019 [cited 2020 Mar 10]. p. 861054. Available from: https://www.biorxiv.org/content/10.1101/861054v1

70. Saunders CT, Wong WSW, Swamy S, Becq J, Murray LJ, Cheetham RK. Strelka: accurate somatic small-variant calling from sequenced tumor-normal sample pairs. Bioinformatics. 2012 Jul 15;28(14):1811–7.

71. Chen X, Schulz-Trieglaff O, Shaw R, Barnes B, Schlesinger F, Källberg M, et al. Manta: rapid detection of structural variants and indels for germline and cancer sequencing applications. Bioinformatics. 2016 Apr 15;32(8):1220–2.

72. Wala JA, Bandopadhayay P, Greenwald NF, O’Rourke R, Sharpe T, Stewart C, et al. SvABA: genome-wide detection of structural variants and indels by local assembly. Genome Res. 2018 Apr;28(4):581–91.

73. Lawrence MS, Stojanov P, Mermel CH, Robinson JT, Garraway LA, Golub TR, et al. Discovery and saturation analysis of cancer genes across 21 tumour types. Nature. 2014 Jan 23;505(7484):495–501.

74. Chen J, Yang H, Teo ASM, Amer LB, Sherbaf FG, Tan CQ, et al. Genomic landscape of lung adenocarcinoma in East Asians. Nat Genet. 2020 Feb;52(2):177–86.

75. Wang S, Tao Z, Wu T, Liu XS. Sigflow: an automated and comprehensive pipeline for cancer genome mutational signature analysis. Bioinformatics. 2021 Jul 12;37(11):1590–2.

76. Wang S, Li H, Song M, Tao Z, Wu T, He Z, et al. Copy number signature analysis tool and its application in prostate cancer reveals distinct mutational processes and clinical outcomes. PLoS Genet. 2021 May;17(5):e1009557.

77. Mayakonda A, Lin DC, Assenov Y, Plass C, Koeffler HP. Maftools: efficient and comprehensive analysis of somatic variants in cancer. Genome Res. 2018 Nov;28(11):1747–56.

78. Gaujoux R, Seoighe C. A flexible R package for nonnegative matrix factorization. BMC Bioinformatics. 2010 Jul 2;11:367.

79. Wickham H. Ggplot2. Wiley Interdiscip Rev Comput Stat. 2011 Mar;3(2):180–5.

80. Kim J, Mouw KW, Polak P, Braunstein LZ, Kamburov A, Kwiatkowski DJ, et al. Somatic ERCC2 mutations are associated with a distinct genomic signature in urothelial tumors. Nat Genet. 2016 Jun;48(6):600–6.

81. Alexandrov LB, Nik-Zainal S, Wedge DC, Campbell PJ, Stratton MR. Deciphering signatures of mutational processes operative in human cancer. Cell Rep. 2013 Jan 31;3(1):246–59.

82. Degasperi A, Amarante TD, Czarnecki J, Shooter S, Zou X, Glodzik D, et al. A practical framework and online tool for mutational signature analyses show inter-tissue variation and driver dependencies. Nat Cancer. 2020 Feb;1(2):249–63.

83. Macintyre G, Goranova TE, De Silva D, Ennis D, Piskorz AM, Eldridge M, et al. Copy number signatures and mutational processes in ovarian carcinoma. Nat Genet. 2018 Sep;50(9):1262–70.

84. Tan VYF, Févotte C. Automatic relevance determination in nonnegative matrix factorization with the β-divergence. IEEE Trans Pattern Anal Mach Intell. 2013 Jul;35(7):1592–605.

85. Bergstrom EN, Huang MN, Mahto U, Barnes M, Stratton MR, Rozen SG, et al. SigProfilerMatrixGenerator: a tool for visualizing and exploring patterns of small mutational events. BMC Genomics. 2019 Aug 30;20(1):685.

86. Ha G, Roth A, Khattra J, Ho J, Yap D, Prentice LM, et al. TITAN: inference of copy number architectures in clonal cell populations from tumor whole-genome sequence data. Genome Res. 2014 Nov;24(11):1881–93.

87. Adalsteinsson VA, Ha G, Freeman SS, Choudhury AD, Stover DG, Parsons HA, et al. Scalable whole-exome sequencing of cell-free DNA reveals high concordance with metastatic tumors. Nat Commun. 2017 Nov 6;8(1):1324.

