## Supplementary figures and images for "Somatic mutation but not aneuploidy differentiates lung cancer in never-smokers and smokers"

### Supplementary Figures

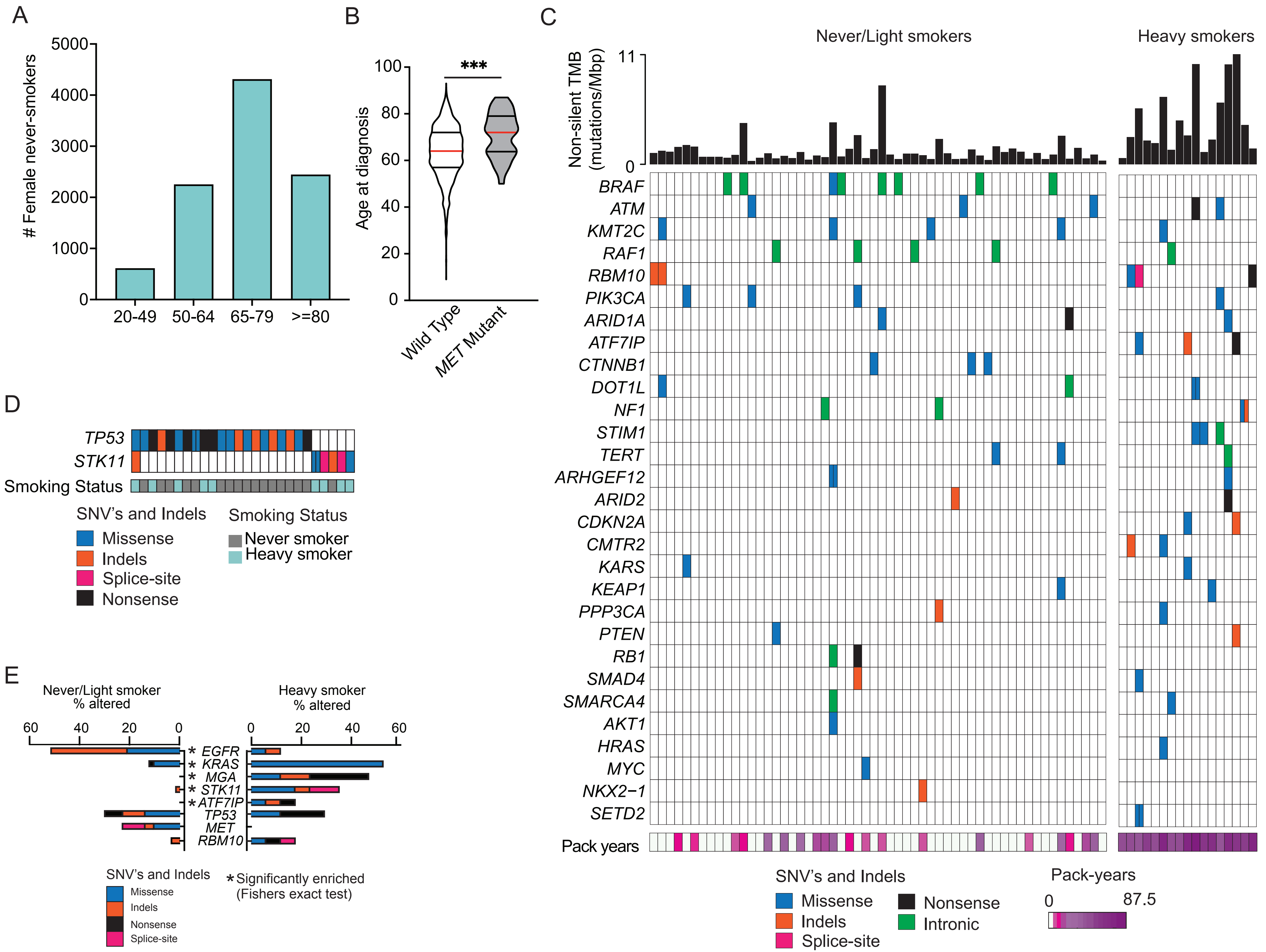

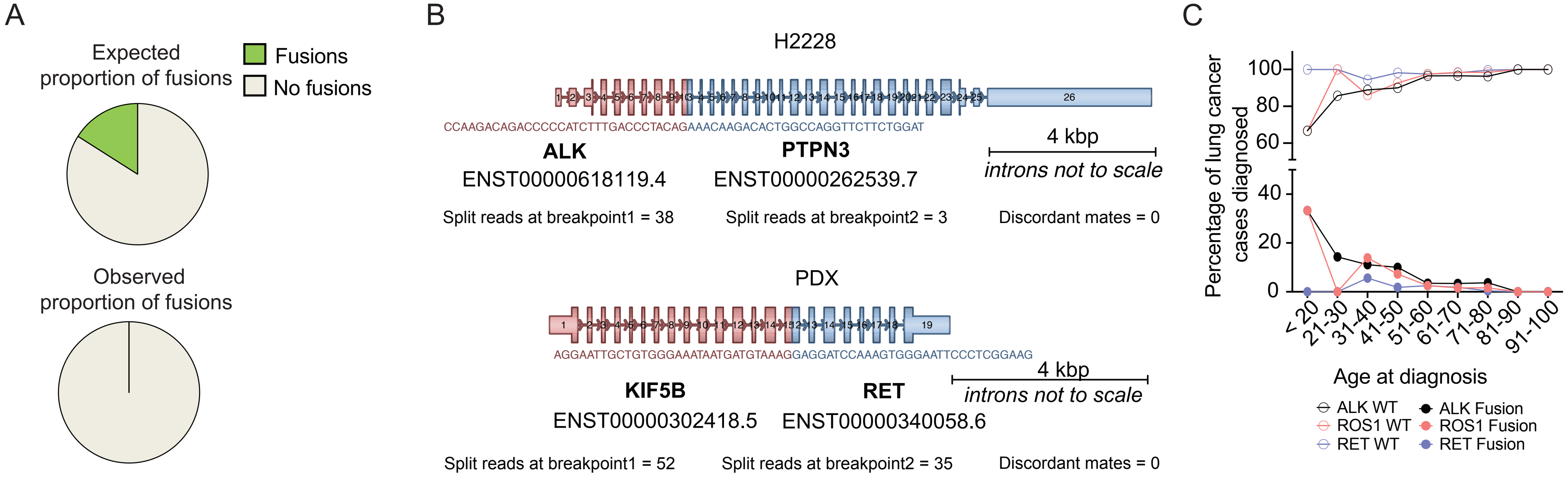

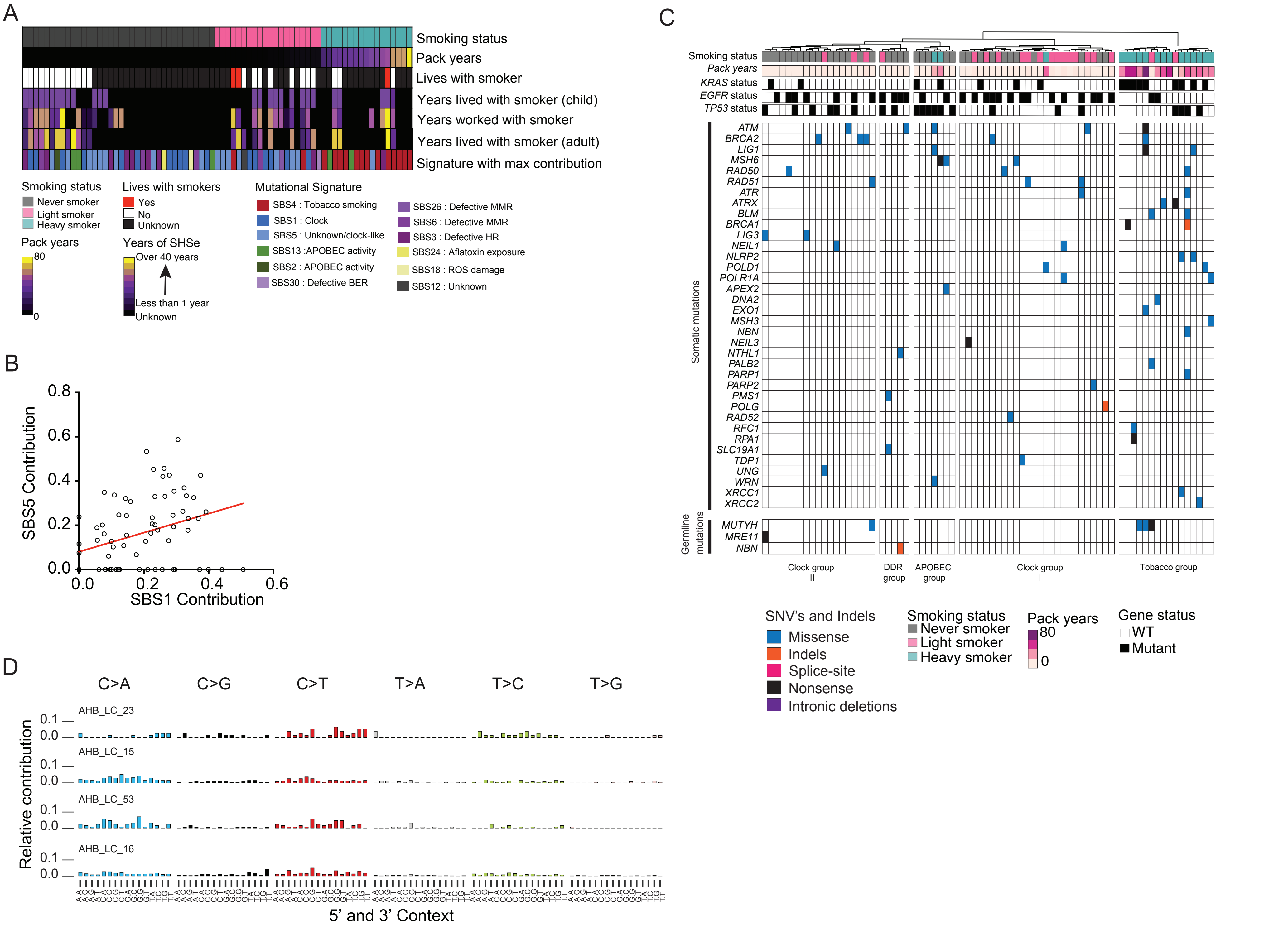

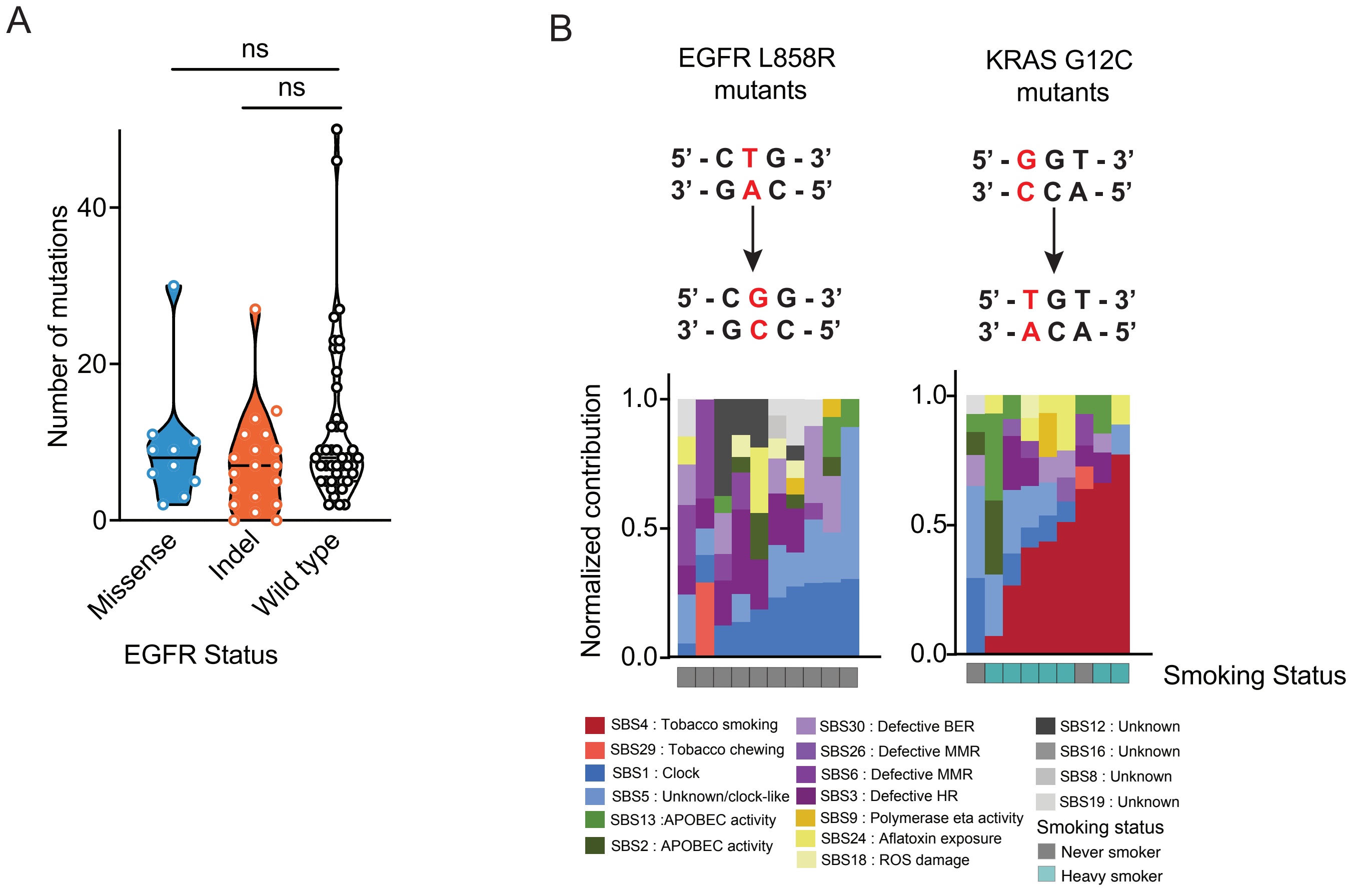

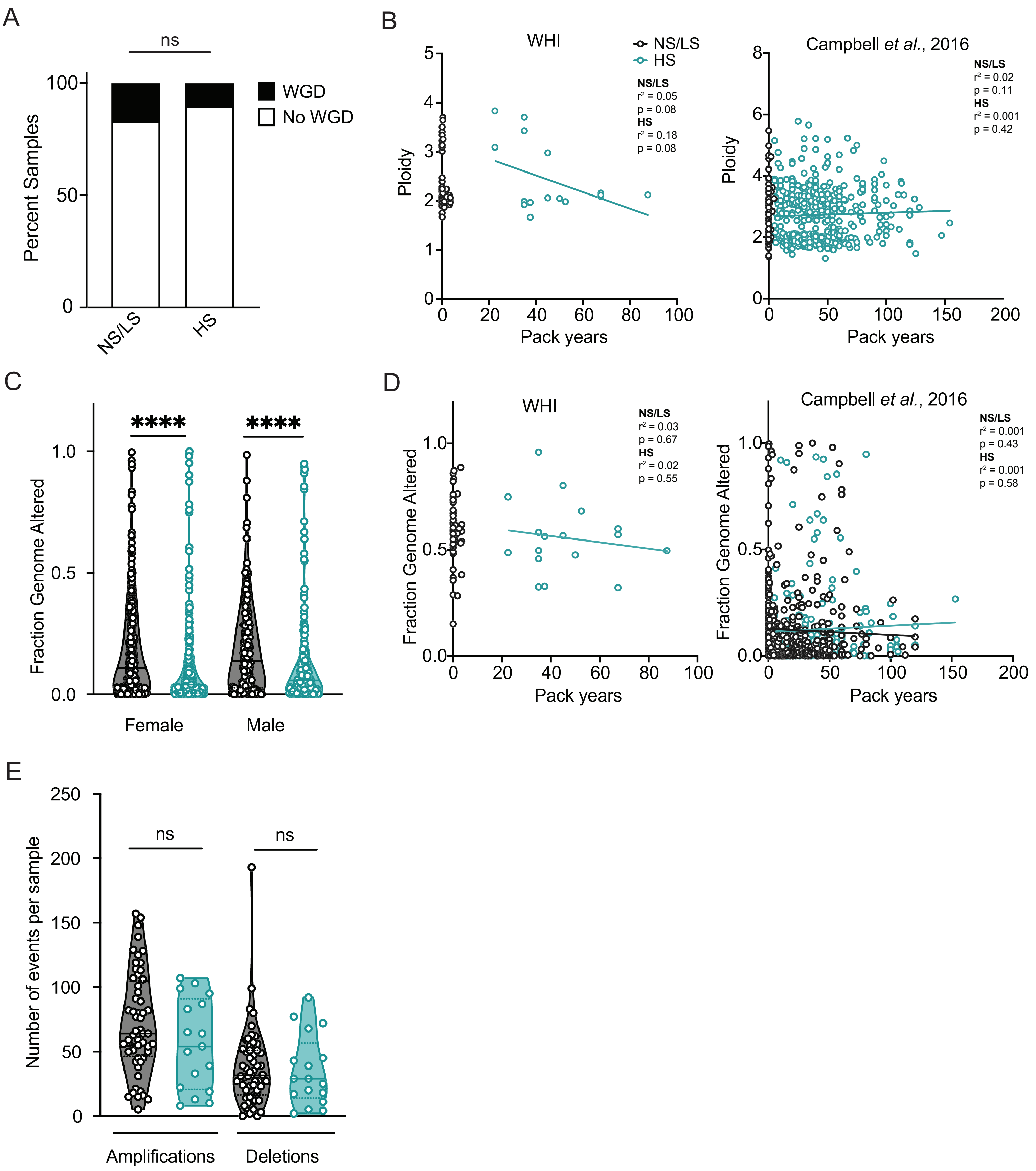
